# On the origin and evolution of microbial mercury methylation

**DOI:** 10.1101/2022.11.21.517362

**Authors:** Heyu Lin, Edmund R. R. Moody, Tom A. Williams, John W. Moreau

**Affiliations:** School of Geographical, Atmospheric and Earth Sciences, The University of Melbourne, Parkville, Victoria 3010, Australia; School of Earth Sciences, University of Bristol, Bristol, BS8 1RJ, United Kingdom; School of Biological Sciences, University of Bristol, Bristol, BS8 1TQ, United Kingdom; School of Geographical and Earth Sciences, University of Glasgow, Glasgow, G12 8RZ, United Kingdom

**Keywords:** mercury, methylmercury, evolution, antimicrobial, *hgc* gene, gene loss, horizontal gene transfer, LUCA

## Abstract

The origin of microbial mercury methylation has long been a mystery. Here we employed genome-resolved phylogenetic analyses to decipher the evolution of the mercury methylating gene, *hgcAB*, constrain the ancestral origin of the *hgc* operon, and explain the distribution of *hgc* in Bacteria and Archaea. We infer the extent to which vertical inheritance and horizontal gene transfer have influenced the evolution of mercury methylators and hypothesize that evolution of this trait bestowed the ability to produce an antimicrobial compound (MeHg^+^) on a potentially resource-limited early Earth. We speculate that, in response, the evolution of MeHg^+^-detoxifying alkylmercury lyase (encoded by *merB*) reduced a selective advantage for mercury methylators and resulted in widespread loss of *hgc* in Bacteria and Archaea.

**Significance:** Neurotoxic methylmercury (MeHg^+^_(aq)_) is synthesized from Hg^II^_(aq)_ in the environment by microorganisms possessing the gene pair *hgcAB*. Our phylogenetic analyses elucidate the origin and evolution of the *hgc* operon, and support a hypothesis that mercury methylation evolved as an antimicrobial production mechanism, possibly from competition for limited resources on the early Earth. We infer from our analyses that *hgc* has been primarily vertically inherited in Bacteria and Archaea, with extensive parallel loss, and note that few taxa possessing *hgc* also possess the gene encoding for MeHg^+^ demethylation, *merB*. Our findings support the interpretation that *merB* evolved as a defense mechanism against the evolution of microbial Hg^II^_(aq)_ methylation.

## Introduction

Microorganisms play key roles in the global mercury cycle. In soils, sediments and natural waters, microbes mediate the enzymatic reductive volatilization of Hg^II^_(aq)_ to Hg^0^_(g)_, the transformation of Hg^II^_(aq)_ to MeHg^+^_(aq)_, the complexation of aqueous Hg^II^_(aq)_ by bacteriogenic bisulfide, and the adsorption/desorption of Hg^II^_(aq)_ on/from biogenic metal ox(yhydrox)ides (e.g., Barkay et al. 2003; Barkay & Gu 2022; Barkay & Wagner□ Döbler 2005; Gilmour et al. 2013; Hsu-Kim et al. 2013; Tebo et al. 2004; Wang et al. 2022). Essentially, microbes control or strongly influence the geochemical speciation of mercury in most natural environments today. Microbial mercury methylation in particular has received much attention because MeHg^+^ is a potent neurotoxin that bioaccumulates across both marine and terrestrial food webs (Kidd et al. 2012).

The origin of microbial mercury methylation has remained a mystery, largely because MeHg^+^ serves no known biological function. Mercury methylation requires Hg^II^_(aq)_, and on the early Earth, volcanoes would have emitted abundant Hg% and some Hg^II^ (Grasby et al. 2019). However, studies have shown that Hg^0^_(g)_ can be efficiently oxidized to Hg^II^_(aq)_ by reactive Cl^-^_(aq)_ and Br^-^_(aq)_ (e.g., Amyot et al. 2005, Seigneur and Lohman, 2008, Obrist et al. 2010, Wang et al. 2014). Volcanos that emitted Hg^II^_(g)_ on the early Earth could also have emitted reactive HCl and BrO (Bobrowski et al. 2003; Guo and Korenaga 2021), providing oxidants to convert Hg^0^_(g)_ to Hg^II^_(aq)_. Recent work supports a dual role for Archean volcanism in Hg^0^_(g)_ and halogen emissions, and subsequent oxidation of Hg^0^_(g)_ to Hg^II^_(aq)_ (Zerkle et al. 2021), as well as the possibility of ephemeral periods of atmospheric oxygenation prior to the Great Oxidation Event at ~2.4 billion years ago (Meixnerova et al. 2021). Following this event, Hg^II^_(aq)_ would have been commonly found as a trace metal in aqueous environments. We assume therefore that Hg^II^_(aq)_ would have been available on the early Earth for methylation upon evolution of the ancestral *hgc* operon. Intriguingly however, the obligately anaerobic and “canonical” Hg methylator, *Pseudodesulfovibrio mercurii*, can apparently oxidize dissolved Hg^0^ directly, and methylate the resulting Hg^II^_(aq)_ (Columbo et al. 2013).

Microbial mercury methylation is currently thought to require the *hgc* operon, most closely related to the ancient carbon monoxide dehydrogenase/acetyl-CoA synthase gene, *cdh*, suggested to have been present in the Last Universal Common Ancestor (“LUCA”; Adam et al. 2018; Parks et al. 2013; Sousa & Martin 2014; Weiss et al. 2016, 2018). The cobalamin-dependent proteins CdhD and CdhE (hereafter “CdhDE”) are present in a highly conserved operon in genomes enabling methyl transfer in the Wood-Ljungdahl (WL) pathway for autotrophic carbon assimilation (Svetlitchnaia et al. 2006). While HgcA is also predicted to be a cobalamin-dependent methyltransferase, it only functions in Hg^II^_(aq)_ methylation and does not seem to take part in the WL pathway, as other components of this pathway are incomplete in characterised *hgcA-carrying* genomes (Date et al. 2019). Similar to *cdh*, *hgc* seems to be broadly distributed across Bacteria and Archaea representing a range of environmental habitats and redox potentials (McDaniel et al. 2020; Capo et al. 2022). Some of these bacteria and archaea have been experimentally confirmed to methylate Hg^II^_(aq)_, while others are still considered as “putative methylators” encoding homologous *hgcAB* sequences that have not yet been experimentally validated. In lieu of cultivated isolates, computational modelling has been employed to test for consistency between putative HgcAB sequences and HgcAB structure and functionality in confirmed methylators (Gionfriddo et al. 2016; Lin et al. 2021).

Many studies of mercury methylation have focused on environmental factors influencing Hg^II^_(aq)_ bioavailability for uptake by methylating cells. In addition, previous efforts to identify the biochemical mechanism of Hg^II^_(aq)_ methylation have attributed the process to 1) accidental enzymatic catalysis of a methyl group transfer to Hg^II^_(aq)_ during either acetyl co-A formation (Choi et al. 1994) or methionine synthesis (Siciliano & Lean 2002), or 2) an as-yet unidentified pathway (Ekstrom et al. 2003). Here, we focus on understanding the more elusive origin and evolution of mercury methylating genes, employing genome-resolved phylogenetic analyses to constrain the ancestry of the *hgc* operon and explain its currently known distribution in Bacteria and Archaea. We assess the extent to which vertical descent and horizontal gene transfer have shaped *hgc* evolution, and propose a hypothesis for the functional origin of microbial mercury methylation as an antimicrobial production mechanism. We also evaluate a potential evolutionary link between *hgc* and *merB*, the latter gene encoding for organomercury lyase (MerB), which lyses the C-Hg bond in MeHg^+^ to release Hg^II^_(aq)_ for reductive volatilization (i.e., detoxification) by MerA (Tezuka and Tonomura 1976).

## Results and Discussion

### *hgc* present in LUCA

A dataset containing 478 protein sequences belonging to the protein family PF03599 was retrieved from UniProt Reference Proteomes database v2022_03 (Chen et al. 2011) at 35% cutoff (RP35) to study the phylogeny of the protein family (**Supplementary Table S1 & S2**). Phylogenetic reconstruction of the protein family PF03599 defined three deep-branching clusters, comprising HgcA, CdhD, and CdhE (**Figure 1A**, for details see **Figure S1**). On the assumption that CdhD- and CdhE-encoding genes were present in LUCA (Adam et al. 2018; Sousa & Martin 2014; Weiss et al. 2018), we deduce that *hgcA* was also likely to be present (i.e., these three protein subfamilies diverged in, or before, LUCA; **Figure 1A**). While robustly rooting single gene trees is challenging, midpoint, Minvar (Mai et al. 2017) and minimal ancestor deviation (MAD) (Tria et al. 2017) rooting approaches all supported a root between CdhD and CdhE+HgcA. In comparison to genes encoding for CdhD and CdhE, which exhibit relatively wide distributions across extant prokaryotes and broadly congruent evolutionary histories, *hgcA* exhibits a more restricted distribution across extant taxa. The latter gene is mainly found in Bacteria, mostly affiliated with Deltaproteobacteria, Firmicutes, and the FCB group, although with a more restricted distribution than in Archaea (**Figure 1A**). Based on the rooted protein tree (**Figure 1B**), HgcA has a longer stem than CdhD and CdhE, which might have resulted from accelerated evolution (e.g., associated with a change in function) or gene loss (or lineage extinction) in former HgcA-encoding clades.

**Figure 1.**
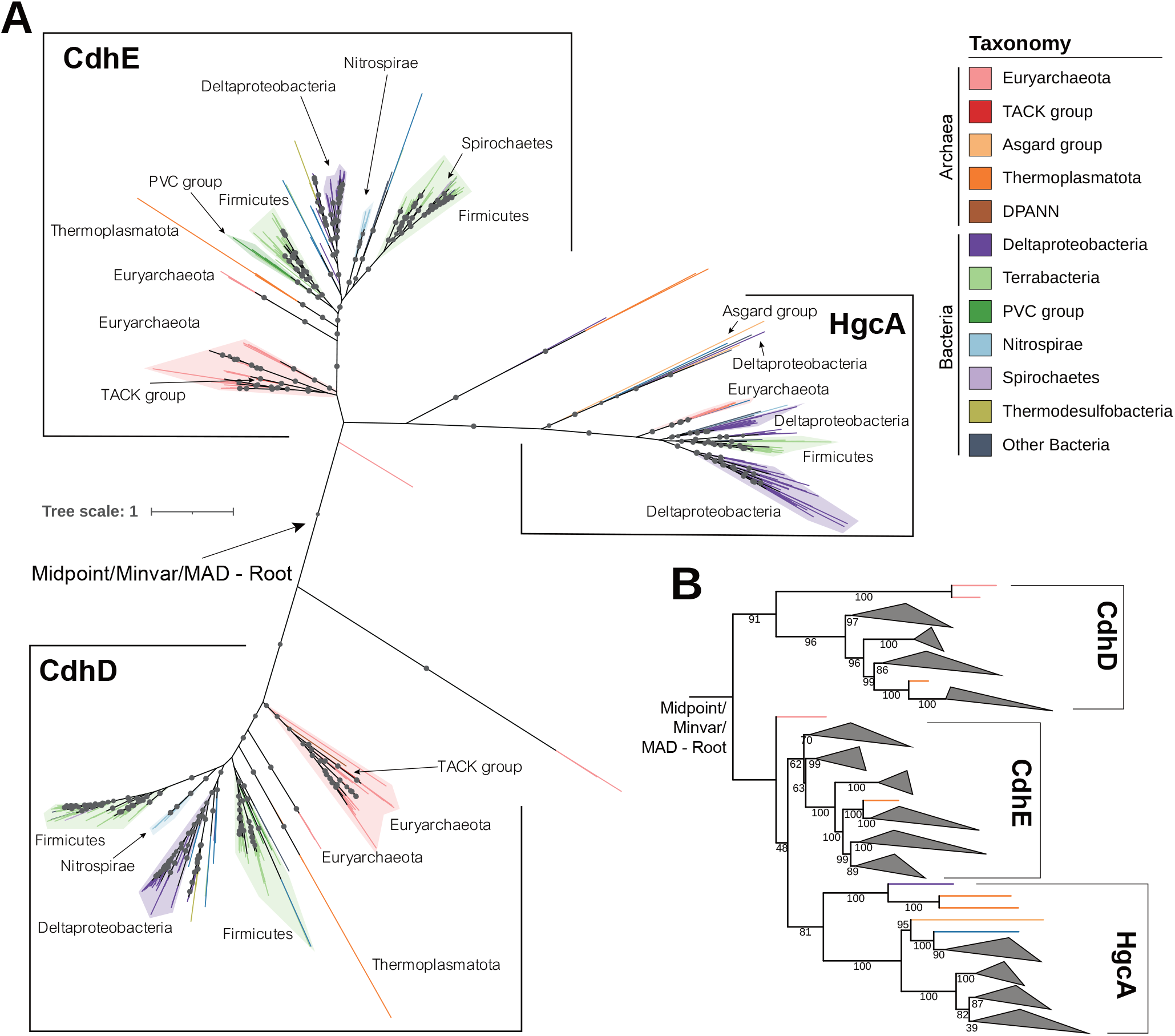
Phylogenetic tree of the protein family PF03599. (A) Unrooted tree of the protein family PF03599. The tree was inferred by using the Maximum Likelihood method under LG+C50+F+R model. This analysis involved 478 amino acid sequences with a total of 2922 positions in the alignments. Different taxonomies are represented by different colors. Ultrafast bootstrap support values were calculated with 1000 replications, and ultrafast bootstrap values > 90% are shown by black dots at the nodes. (B) PF03599 tree rooted by the midpoint. Clades whose average branch length distance to their leaves are below 1.5 are collapsed with iTOL for better visualization.

### *hgcA* evolved primarily vertically but with extensive loss

In our phylogenetic analysis, to increase the number and taxonomic coverage of targets, we further enlarged the sample size by retrieving HgcA homologs in UniProt Reference Proteomes database v2022_03 at 75% cutoff (RP75). Two other datasets, including one containing 700 representative prokaryotic proteomes constructed by Moody et al. (2022) and another containing several novel *hgc*-carriers published by Lin et al. (2021), were retrieved and incorporated into the RP75 dataset. After removing redundancies and including sequences from new phyla the number of HgcA sequences in the dataset was increased from 76 to 169 (**Supplementary Table S1**). The resulting HgcA tree was rooted according to the topology of the HgcA subtree in **Figure 1**.

Our analysis suggests that *hgcA* genes are separated into two well-supported clades with high bootstrap support, as shown in **Figure 1A** and **Figure 2** (for details see **Figure S2**). These two clades correspond to a difference in gene structure: the smaller clade comprises genes in which *hgcA* and *hgcB* became fused to form *hgcAB*, while the larger contains single subunit (*hgcA*) genes. Several Asgard archaea, including some Thorarchaeota and Lokiarchaeota, form a clade in the HgcAB subtree, suggesting that gene fusion occurred prior to the radiation of Asgard phyla. The fused *hgcAB* of *Pyrococcus furiosus* has been experimentally shown not to methylate Hg^II^_(aq)_ (Podar et al. 2015), but the functionality of other fused-*hgcAB* genes, as well as other members of this clade, is yet to be confirmed. For example, *Nitrospina-like hgcA* sequences have been reported as the dominant *hgcA* homologs in many marine environments (Gionfriddo et al. 2016; Tada et al. 2020; Villar et al. 2020; Tada et al. 2021). However, the functionality of these sequences has not yet been experimentally confirmed, as no known *hgc*+ members of this genus are currently held in isolation. Also, the alphaproteobacterium *Breoghania sp*. L-A4 is the only aerobic *hgcA+* (fused-*hgcAB*+) microorganism found in this analysis, and its methylation capacity under aerobic conditions also remains untested.

**Figure 2.**
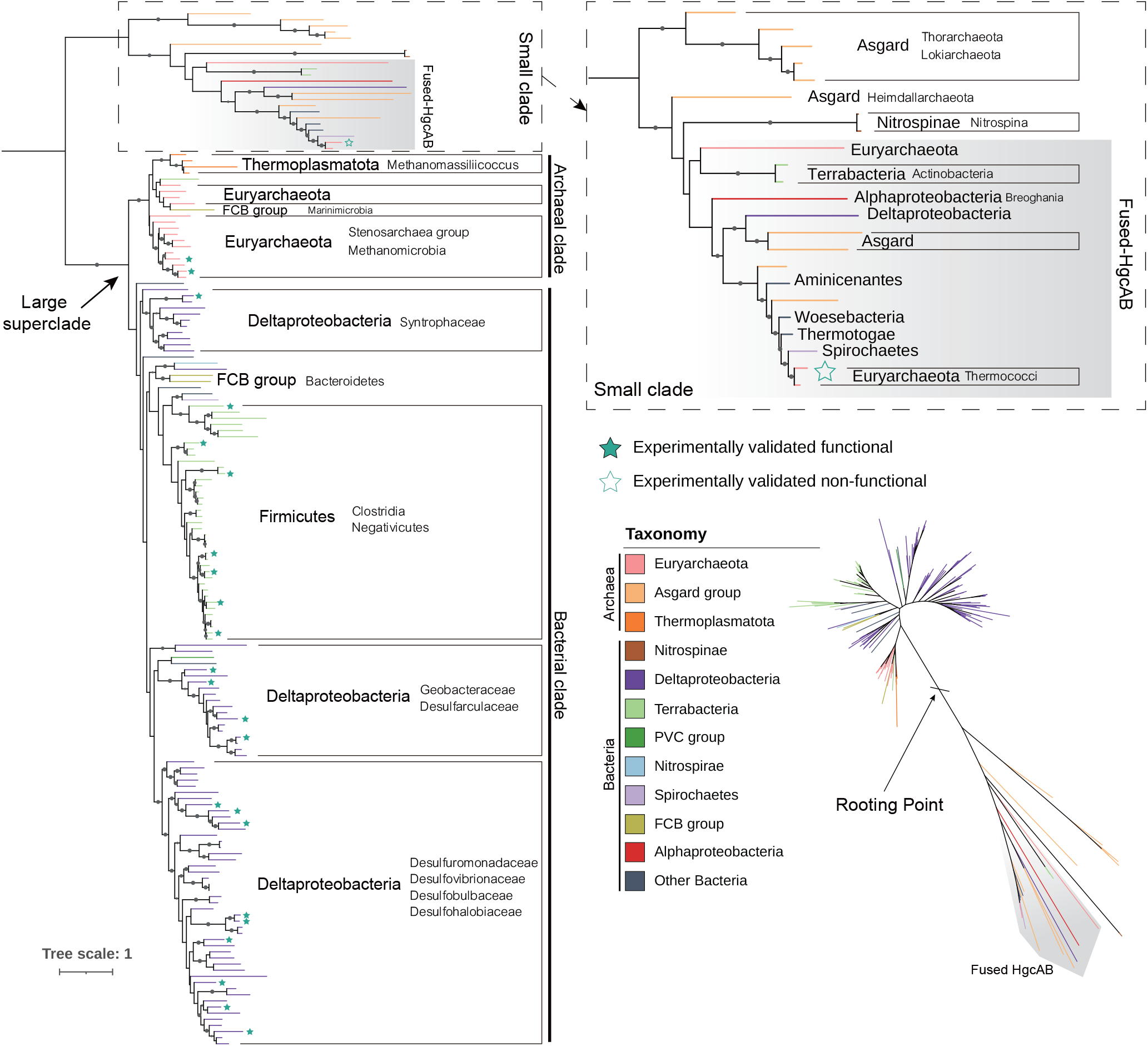
Phylogenetic tree of HgcA proteins. The tree was inferred by using the Maximum Likelihood method under LG+C60+F+G model. This analysis involved alignment of 169 amino acid sequences with a total of 493 positions. Different taxonomies are represented by different colors. Ultrabootstrap support values were calculated with 1000 replications, and ultrabootstrap values > 90% are shown by black dots at the nodes. Experimentally validated functional and non-functional HgcA sequences are labelled by solid and hollow stars. Fused-HgcAB sequences are indicated by a grey background.

By contrast, the majority of (non-fused) HgcA sequences form a clade which broadly follows the universal tree (**Figure 2**). In particular, the deepest split lies between archaeal and bacterial HgcA homologs. Taken together with the sister relationship of HgcA to CdhD/E, this finding supports an origin of HgcA in or before LUCA. However, the taxonomic distribution of HgcA in extant Bacteria is patchy (being found mainly in Firmicutes, Deltaproteobacteria, and the FCB group). Therefore, if HgcA was indeed present in LUCA, and its gene tree traces the divergence of Archaea and Bacteria, it must have been lost subsequently in the other bacterial lineages. While it is undoubtably difficult to resolve the evolutionary history of ancient single genes with certainty, it is interesting that an ancient origin for HgcA cannot be excluded based on the inferred tree.

Gene loss is a major force driving microbial genome evolution (Bolotin & Hershberg 2016), reflecting environmental change and/or adaptation (Koskiniemi et al. 2012). So far, *hgcA*+ methylators have primarily been reported for anoxic or suboxic environments (Capo et al. 2022; Lin et al. 2021; Parks et al. 2013). Changes in redox potential that could inhibit Hg^II^_(aq)_ methylation may have presented a selective environmental pressure to lose *hgcA*. In addition, long branches of the HgcA subtree, in comparison to CdhD/E, also suggest a weaker natural selection for gene maintenance, and greater likelihood of loss through genetic drift, a common scenario observed for many microorganisms (Bolotin & Hershberg 2016).

A few putative HGT events were inferred for the larger clade of the HgcA tree, e.g., Marinimicrobia-HgcA clustered with Euryarchaeota-HgcA in the archaeal cluster, with 98% coverage and 56% identity by blastp against HgcA from Theionarchaea archaeon DG-70-1, suggesting a possible lateral acquisition of *hgcA*. This inference is consistent with that of McDaniel et al. (2020) who also suggested HGT played an important role in HgcA evolution. Our study differs from that of McDaniel et al. in suggesting that *hgcA* may date back to the LUCA; this inference is based on the nesting of the HgcA subtree within the broader diversity of the Cdh family (**Figure 1B**), an aspect of the family’s evolutionary history not considered in that analysis, as well as the recovery in our analyses of a deep split between archaeal and bacterial sequences in the hgcA subtree, notwithstanding some HGT.

### *hgcA* and *hgcB* co-evolved

The gene *hgcB* is nearly always located immediately downstream of *hgcA*, with rare cases of being one gene apart (Gilmour et al. 2018). A lack of either *hgcA* or *hgcB* in a genome is thought to render the microbial host incapable of Hg^II^_(aq)_ methylation (Parks et al. 2013). Therefore, the gene pair *hgcAB* likely evolved together as a conserved operon. Although the HgcB tree was poorly resolved due to short sequence length (95 amino acids on average), the overall topologies of the HgcB tree and the HgcA tree were congruent, supporting this inference (**Figure 3**, for details see **Figure S3**). Nevertheless, several *hgcA+* genomes did not carry neighbouring *hgcB* genes, including all *Nitrospina* and a few Deltaproteobacteria and Firmicutes, potentially because of gene loss during evolution or incomplete transfer events (i.e., only *hgcA* genes were acquired during the HGT events). Another exception involves a complete *hgcB* gene present downstream of the fused-*hgcAB* gene in the genome of the deltaproteobacterial endosymbiont, Delta1. This complete HgcB protein had the longest branch length of the HgcB tree and was located in a sister cluster of the ‘HgcB tail’ associated with the fused-*hgcAB* from the same host, suggesting a possible lateral acquisition of *hgcB* from closely related taxa.

**Figure 3.**
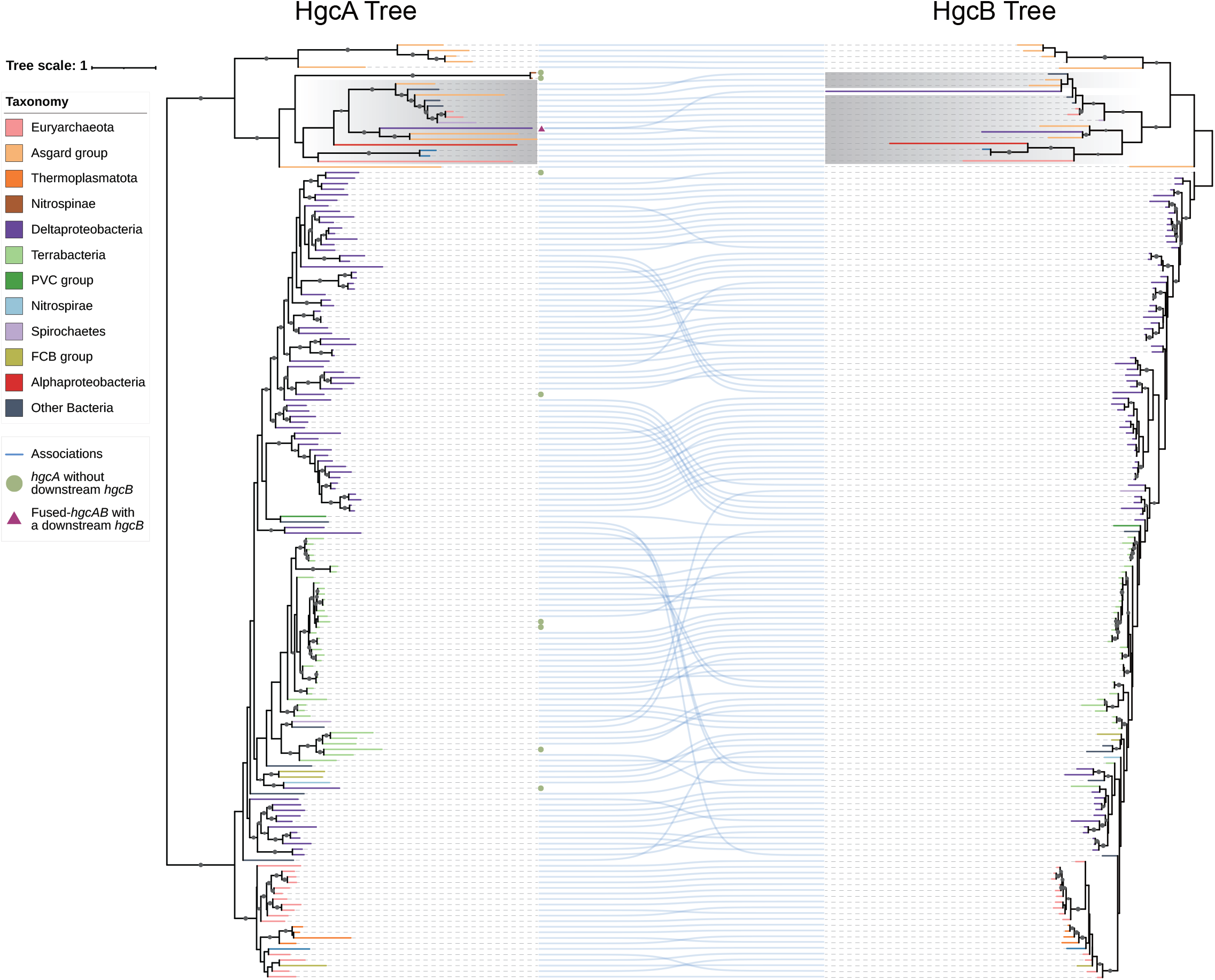
Co-phylogenetic tree of HgcA and HgcB. The phylogeny of HgcA shown on the left is the same as Figure 2. The phylogeny of HgcB shown on the right was inferred by using the Maximum Likelihood method under LG+C60+F+G model. Lines between the two trees connect HgcA (left) and HgcB (right) proteins from the same proteome, illustrating the degree of congruency in their respective phylogenetic associations. This analysis involved alignment of 169 amino acid sequences with a total of 322 positions. Taxonomies of the HgcB sequences are represented in the same colors as shown in the HgcA phylogeny. Ultrabootstrap support values were calculated with 1000 replications, and ultrabootstrap values > 90% are shown by black dots at the nodes. Gray shaded blocks describe fused-HgcAB and ‘HgcB tail’ genes in the two trees, respectively. The green dots at the tip of the HgcA tree represent the corresponding *hgcA* without downstream *hgcB*, and the red triangle represents the corresponding fused-*hgcAB* with another *hgcB* downstream.

### Phylogenetic distribution of *merB* and its relationship to *hgc*

Microbial detoxification mechanisms for Hg^II^_(aq)_ and MeHg^+^_(aq)_, including the enzymes MerA and MerB, respectively, have been well described (c.f., Robinson and Tuovinen 1984, Barkay et al. 2010, Barkay & Gu 2022). Recent phylogenetic analyses have revealed a thermophilic and archaeal origin for the *mer* operon, acquired later by bacteria through multiple independent transfer events (Christakis et al. 2021) reflective of the environmental advantage of mercury resistance genes (Boyd & Barkay 2012). In total, 225 MerB sequences were retrieved from the RP35 dataset (**Supplementary Table S2**). MerB homologs were distributed across Archaea and Bacteria (**Figure 4**, for details see **Figure S4**), with most representatives found in Terrabacteria (including Actinobacteria, Firmicutes, and Chloroflexi) and Proteobacteria (including Alphaproteobacteria, Betaproteobacteria, and Gammaproteobacteria). In contrast, only a small number of archaea, including a few euryarchaeotes and one Thaumarchaeon, were found to encode MerB. These archaeal MerB homologs were placed within bacterial MerB clades and were not monophyletic, consistent with the interpretation of recruitment of *merB* to the *mer* operon from a mesophilic ancestor (Christakis et al. 2021), which we infer involved HGT from Bacteria to Archaea.

**Figure 4.**
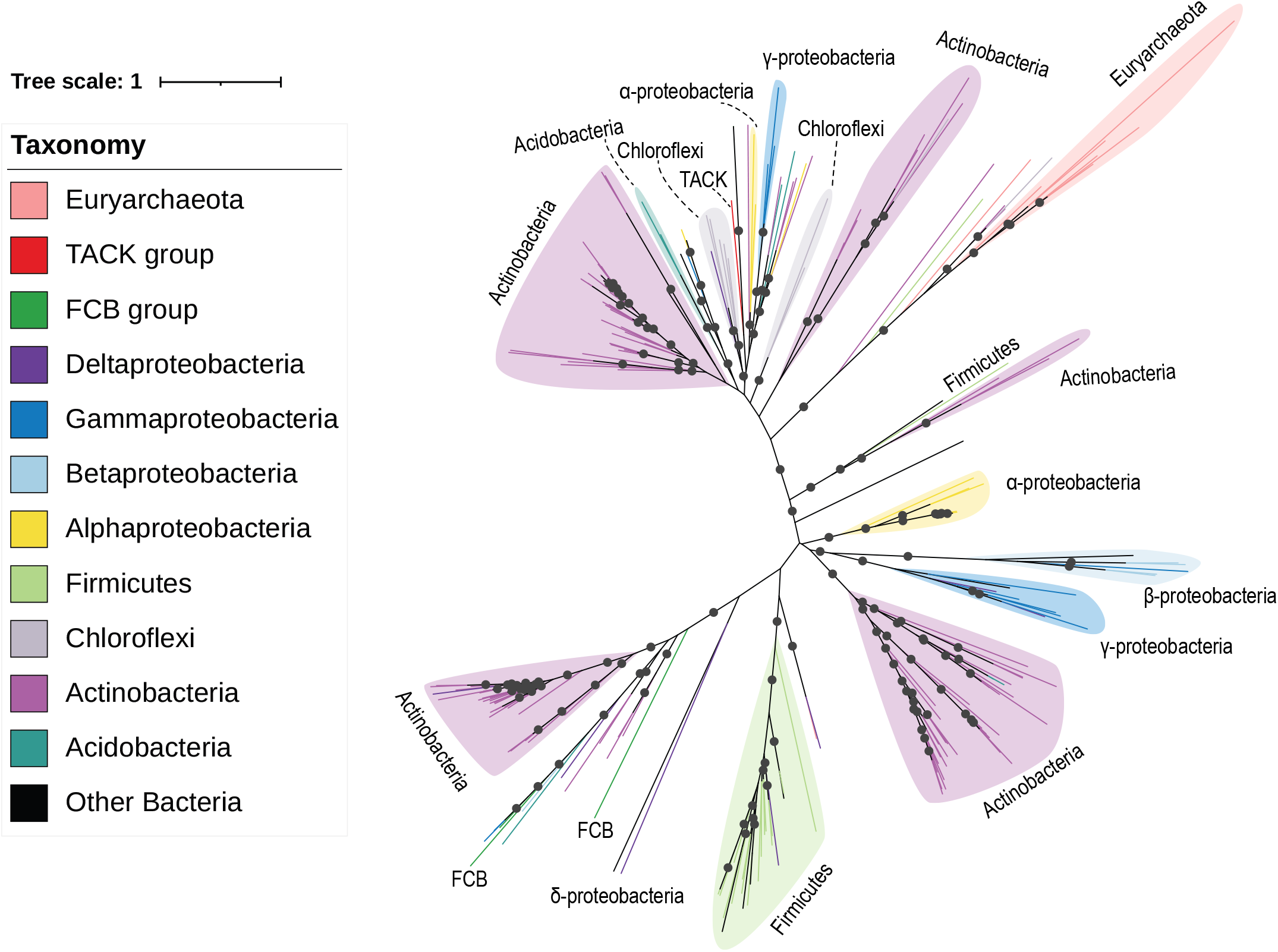
Phylogenetic tree of MerB proteins. The tree was inferred by using the Maximum Likelihood method under LG+C40+F+G model. This analysis involved alignment of 223 amino acid sequences with a total of 1448 positions. Different taxonomies are represented by different colors. Ultrabootstrap support values were calculated with 1000 replications, and ultrabootstrap values > 90% are shown by black dots at the nodes.

We mapped the presence/absence of *merB* and *hgc* genes onto the species tree (**Figure 5**, for details see **Figure S5**) to investigate their phylogenetic relationships. Intriguingly, although the distribution of MerB homologs overlapped with HgcAB homologs, few genomes encode *both* MerB and HgcAB, signifying that mecury methylators generally cannot also demethylate MeHg^+^. This apparent polarization in gene distribution suggests a functional conflict between these two genes (i.e., there is utility in encoding for one or the other function, but not both). Only a few genomes were found to encode for both MerB and HgcAB; this dual functionality in *Citrifermentans bemidjiense* Bem was previously recognized and discussed (Lu et al. 2016). The phylogenetic distribution of these opposing enzymes (**Figure 4 & Figure 5**) suggests that HgcAB was the earlier of the two to evolve, with MerB originating more recently and radiating via horizontal gene transfer.

**Figure 5.**
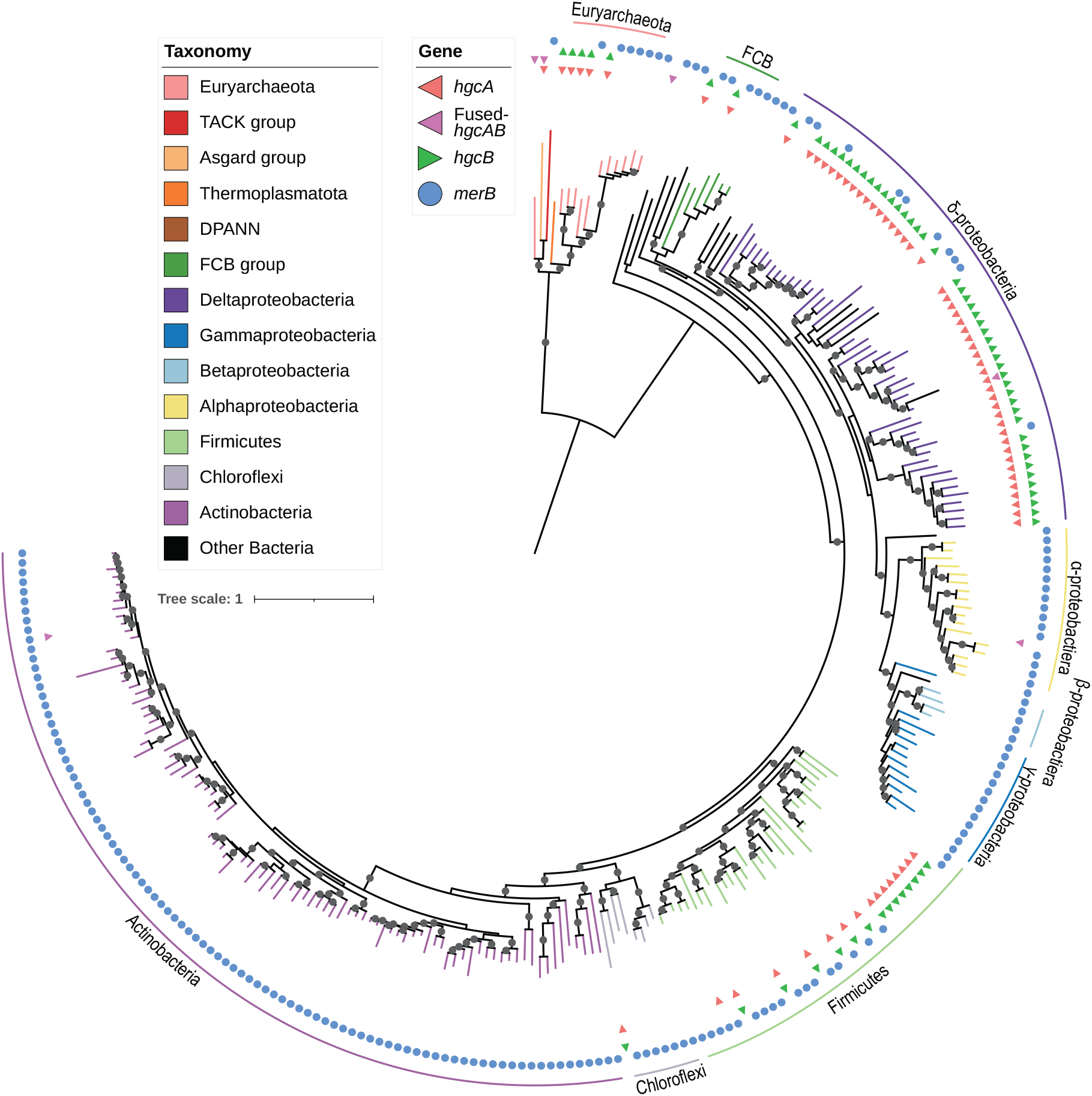
Species tree of *hgcA+* and *merB+* genomes in the RP35 dataset. The tree was inferred from a concatenate of 27 marker genes derived from Moody et al. (2022) by using the Maximum Likelihood method under LG+F+R10 model. This analysis involved alignment of 251 amino acid sequences with a total of 8024 positions. Different taxonomies are represented by different colors. The presence of genes *hgcA*/fused *hgcAB*/*hgcB*/*merB* in the genomes were represented by different symbols and colors in the outer circle. Ultrabootstrap support values were calculated with 1000 replications, and ultrabootstrap values > 90% are shown by black dots at the nodes.

### Mercury methylation as antimicrobial synthesis?

The reason why microorganisms evolved the ability to methylate Hg^II^ to MeHg^+^ has long been a mystery. Previous studies proposed that this process might reduce susceptibility to Hg^II^_(aq)_ toxicity (Trevors 1986). But in fact, the ability to produce MeHg^+^_(aq)_ has been shown *not* to confer Hg^II^_(aq)_ resistance in methylating microbes (Gilmour et al. 2011). Previous studies have suggested that MeHg^+^_(aq)_ is more toxic than Hg^II^_(aq)_ to some microbes (Jonas et al. 1984; Gilmour et al. 2011), and we hypothesise here that Hg^II^_(aq)_ methylation may have evolved effectively as an early antimicrobial against microorganisms without the ability to metabolize/detoxify this organometallic compound. We note that, in seawater, the higher proportion of lipophilic MeHg^+^_(aq)_ (speciated as MeHgCl(aq)) over Hg^II^_(aq)_ (as HgCl2(aq)) renders the former a more effective toxicant to microbial cells (Mason et al. 1995). Microorganisms have evolved a wide range of strategies to compete against other microbes for limited resources (Granato et al. 2019), and our hypothesis that MeHg^+^_(aq)_ effectively originated as an antimicrobial mirrors a similar interpretation for *arsM* encoding for arsenic methylation (Li et al. 2021). As the *hgcAB* gene pair likely evolved in LUCA (**Figure 1A),** Hg^II^_(aq)_ methylation could have stabilised to persist as an early form of antimicrobial production.

As a defense against MeHg^+^_(aq)_, other microorganisms evolved alkylmercury lyase (MerB) encoded by *merB*, with Hg^II^_(aq)_ toxicity mitigated via other detoxification systems, e.g., MerA (Boyd & Barkay 2012), iron-coupled redox reactions (Liu & Wiatrowski 2018), abiotic reduction (Gu et al. 2011) or other mechanisms (Christakis et al. 2021). Once *merB* evolved however, *hgcAB* may no longer have offered a selective advantage, and therefore underwent extensive loss during vertical evolution. Furthermore, we suggest that bacteria carrying other MeHg^+^_(aq)_ detoxification mechanisms might not need to also carry *merB*. For example, MerB homologs are rarely found in sulfate-reducing bacteria (SRB), as SRB produce H2S as a metabolite that reacts with MeHg^+^_(aq)_ to form dimethylmercury (DMHg) which volatilises efficiently from the cell (Baldi et al. 1993). Jonsson et al. (2016) also found DMHg production resulting from reaction of MeHg^+^_(aq)_ with poorly crystalline iron sulfides; these phases are typically precipitated indirectly by the metabolism of SRB. Other studies have observed that MeHg^+^_(aq)_ is somehow efficiently exported from methylating cells, thus reducing toxicity to the methylator (Graham et al. 2012; Lin et al. 2015). We recognise, however, that the mechanisms underpinning MeHg^+^_(aq)_ export have not yet been elucidated. Interestingly, extracellular thiol compounds, such as cysteine, were found to facilitate export and desorption of MeHg^+^_(aq)_ (Lin et al. 2015), possibly due to competitive binding of thiols to receptors on MeHg^+^_(aq)_ transporters. To our knowledge, only two proteins encoded by *mer* operons, MerC (Sone et al. 2017) and MerE (Sone et al. 2013), have been reported as potential MeHg^+^_(aq)_ transporters, but neither of them is carried by Hg^II^_(aq)_ methylators.

Our study has revealed an ancient origin for microbial mercury methylation, evolving from LUCA to radiate extensively throughout the tree of life both vertically, albeit with extensive loss, and to a lesser extent horizontally. We hypothesise that early mercury methylating microorganisms may have innovatively transformed a ubiquitous aqueous trace metal, Hg^II^_(aq)_, into a highly effective antimicrobial compound. The dual ability of sulfate reducing bacteria to methylate mercury and excrete sulfide provides serendipitous protection against MeHg^+^_(aq)_ toxicity. Methylmercury degradation can also be abiotically photo-catalysed, so it is unlikely that shallow aqueous environments where UV light could penetrate would have allowed for toxic build-up of MeHg^+^_(aq)_. We speculate that anoxic sediments where volcanogenic mercury and either geogenic or bacteriogenic sulfide were co-present (such as seafloor sediments near hydrothermal vents) may have provided an ideal biogeochemical setting for the evolution of microbial mercury methylation. Prior to the evolution of genomic countermeasures such as *merB*, MeHg^+^_(aq)_ concentrations in some natural environments could have increased to toxic levels beyond those typically found today, where they are constrained by both biotic and abiotic demethylation mechanisms. We conclude that microbial Hg^II^_(aq)_ methylation in extant microorganisms represents the genomic vestige of an early geosphere-biosphere interaction with major ramifications for Earth’s mercury cycle.

## Materials and Methods

### Dataset construction

UniProt Reference Proteomes v2022_03 (https://proteininformationresource.org/rps/) at 35% cutoff (RP35) and 75% cut-off (RP75) were used to establish two datasets. UniProt Reference Proteomes is a database containing representative proteomes selected from a large proteome database with more than 30,000 proteomes from UniProtKB (Chen et al. 2011). Every proteome in the database is comprised of all coding sequences translated from a whole genome sequence. The RP35 dataset contained 5,535 proteomes and was used to study the phylogeny of protein family PF03599 (CdhD, CdhE and HgcA) and PF03243 (MerB). The RP75 dataset contained 17,551 proteomes and was used to study the phylogeny of HgcA individually. In order to enlarge the diversity and sample size of HgcA sequences, two other datasets, including one containing 700 representative prokaryotic proteomes constructed by Moody et al. (2022) and another containing several novel *hgc*-carriers published by Lin et al. (2021), were retrieved and incorporated into the RP75 dataset, followed by removal of any redundancies. HgcAB proteins selected for this study including their metadata are listed in Supplementary Table S1; Other proteins including CdhD, CdhE, and MerB used in this study are listed in Supplementary Table S2.

### Determination of genes of interest

Protein sequences belonging to protein families PF03599 and PF03243 from both datasets were determined according to annotation by UniProt and also confirmed with hmmsearch v3.1 (Eddy 1998) against curated HMM models provided by InterPro (https://www.ebi.ac.uk/interpro/entry/pfam). HgcA sequences were extracted from the PF03599 family using hmmsearch against the Hg-MATE database (Capo et al. 2022) and further determined by the conserved motif N(V/I)WC(A/S). CdhD and CdhE sequences were extracted from the PF03599 family using blastp v2.11.0 against the experimentally validated CdhD (Q57577) and CdhE (Q57576) sequences from *Methanocaldococcus jannaschii* JAL-1, respectively. HgcB sequences were determined by searching for 4Fe4S proteins encoded by genes adjacent to *hgcA* genes (two open reading frames on either side of *hgcA*) in the genome. MerB sequences were further confirmed from the PF03243 protein family by hmmsearch against the MerB database constructed by Christakis et al. (2021).

### Reconstruction of phylogenetic trees

Protein sequences belonging to the PF03599 protein family and included in the RP35 dataset were aligned using MAFFT-linsi v7.453 (Katoh & Standley 2013), resulting in an alignment containing 478 sequences with 2922 columns. The alignment was used to build a maximum likelihood (ML) tree using IQ-TREE v2.0.3 (Schmidt et al. 2014) under the best-fitting model of LG+C50+F+R. Branch supports were estimated with 1000 ultrafast bootstrap (Hoang et al. 2018) replicates (**Figure 1**) The unrooted tree was rooted using three methods: Midpoint-rooting (Swofford et al. 1996), Minimum Variance (MinVar, Mai et al. 2017), and Minimal Ancestor Deviation (MAD, Tria et al. 2017), respectively. Similar to the methods described above, all trees in this study were built using IQ-TREE v2.0.3 (Schmidt et al. 2014) with 1000 ultrafast bootstrap (Hoang et al. 2018) replications. Protein sequences were aligned using MAFFT (L-INS-i) v7.453 (Katoh & Standley 2013).

To mitigate potential contamination of HgcB tail sequences in the HgcA alignment, the fused HgcAB proteins were aligned separately and the ‘HgcB tail’ (last 79 positions) was manually removed. The resulting ‘HgcA head’ of the fused HgcAB proteins was aligned with other HgcA sequences. Finally, aligned ‘HgcB tail’ sequences were subsequently added back to the alignment. The first 327 positions of the alignment were poorly aligned, therefore manually removed. The final alignment contained 169 sequences with 493 columns and was used to infer a ML tree under the LG+C60+F+G model (**Figure 2**). The tree was rerooted between the two major groups of HgcA sequences according to the above PF03599 protein family tree.

HgcB sequences and ‘HgcB tail’ sequences from the fused HgcAB proteins were aligned and used to infer a ML tree using a similar method as the PF03599 protein family tree described above under the LG+C60+F+G model. Phylogenetic congruence between the HgcA and the HgcB sequences was inferred and visualized using cophylo implemented in the R package phytools v1.0-3 (Revell 2012) based on the topology of the HgcA and the HgcB ML trees described above. Nodes of both HgcA and HgcB trees were allowed to be rotated by the program to optimize vertical matching of tips (**Figure 3**).

In total, 223 MerB sequences were predicted from proteomes in the RP35 dataset. These sequences were aligned and used to build a ML tree under the best-fit protein model LG+C40+F+G chosen according to BIC **(Figure 4)**.

A species tree of *hgc+* and *merB+* proteomes (251 proteomes in total) from the RP35 dataset was reconstructed based on the 27 marker genes proposed by Moody et al. (2022). 27 HMM profiles were created individually based on marker gene alignments and concatenated using HMMER v3.1. The 27 marker homologs in the 251 proteomes were extracted by hmmsearch against the concatenated hmm profile with an E-value of 1e-10. The 27 marker homologs were then aligned respectively and concatenated. Poorly aligned regions were trimmed using trimAl v1.2 (Silla-Martínez et al. 2009) with parameters “-resoverlap 0.55 -seqoverlap 60 -automated1”. The resulting supermatrix (8024 sites) was used to infer a ML tree under the model of LG+F+R10 (**Figure 5**). All phylogenetic trees described above were visualized using iTOL v6 (Letunic & Bork 2021) and refined with Adobe Illustrator (Adobe Systems Inc., San Jose, CA, USA).

## Data availability

All data generated in this study including amino acid alignments and phylogenetic trees are deposited in Figshare: https://doi.org/10.6084/m9.figshare.21428523

## Acknowledgments

We thank Yao-ban Chan and Qiuyi Li at The University of Melbourne for valuable discussions and advice in the early stage of this study. We also acknowledge helpful discussions with Caitlin Gionfriddo, Ben Peterson, and Eric Capo. This publication was made possible in part through support from a Strategic Australian Postgraduate Award (PhD scholarship) to H.L. from the Environmental Microbiology Research Initiative (EMRI) at The University of Melbourne (J.W.M.) and laboratory start-up funding from the University of Glasgow to J.W.M., the Moore Foundation (https://doi.org/10.37807/GBMF9741 to T.A.W.), the Royal Society (URF\R\201024) to T.A.W., and the John Templeton Foundation (62220 to T.A.W. and E.R.R.M.). The opinions expressed in this publication are those of the author(s) and do not necessarily reflect the views of the John Templeton Foundation.

