## Supplemental Figure S1-S5 for "On the origin and evolution of microbial mercury methylation"

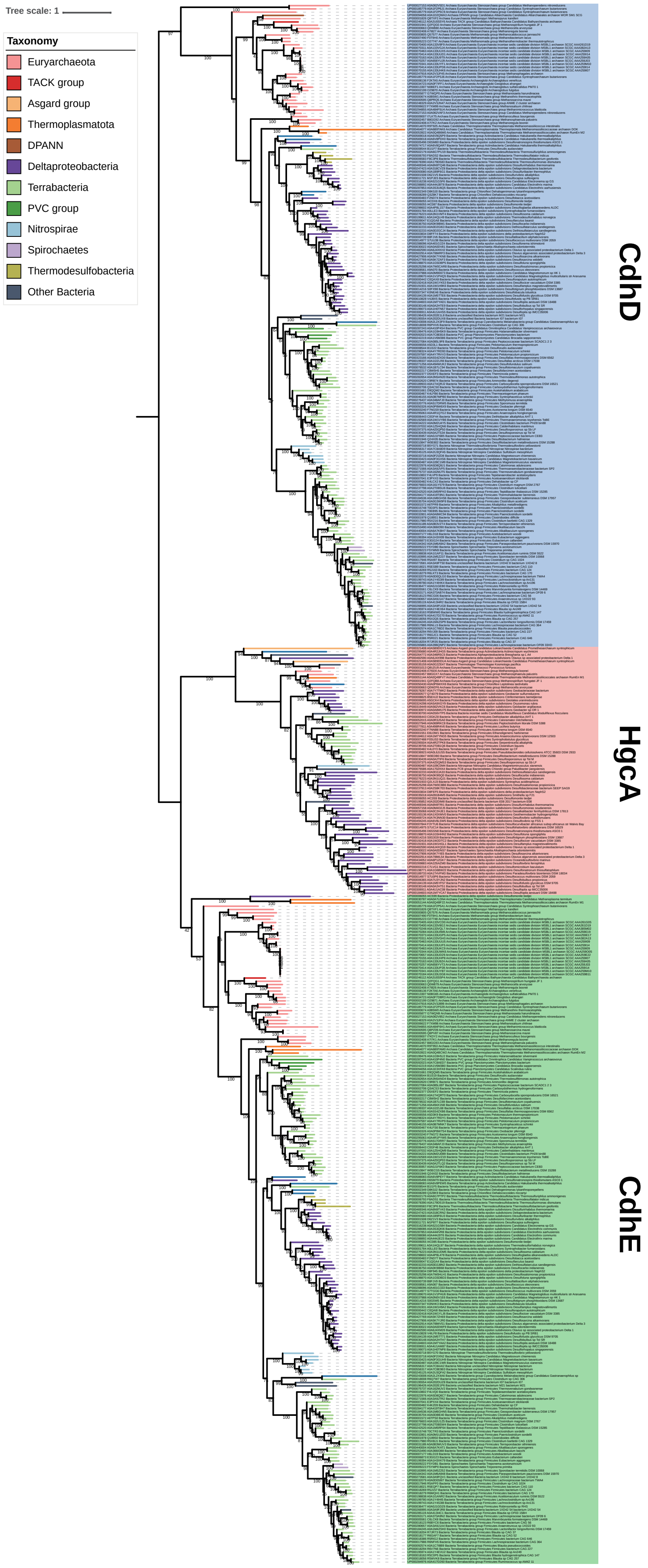


**Figure S1. Phylogenetic tree of the protein family PF03599.** The tree was inferred by using the Maximum Likelihood method under LG+C50+F+R model and rooted by the midpoint. This analysis involved 478 amino acid sequences with a total of 2922 positions in the alignments. Different taxonomies are represented by different colors. Ultrafast bootstrap support values shown at nodes were calculated with 1000 replications.

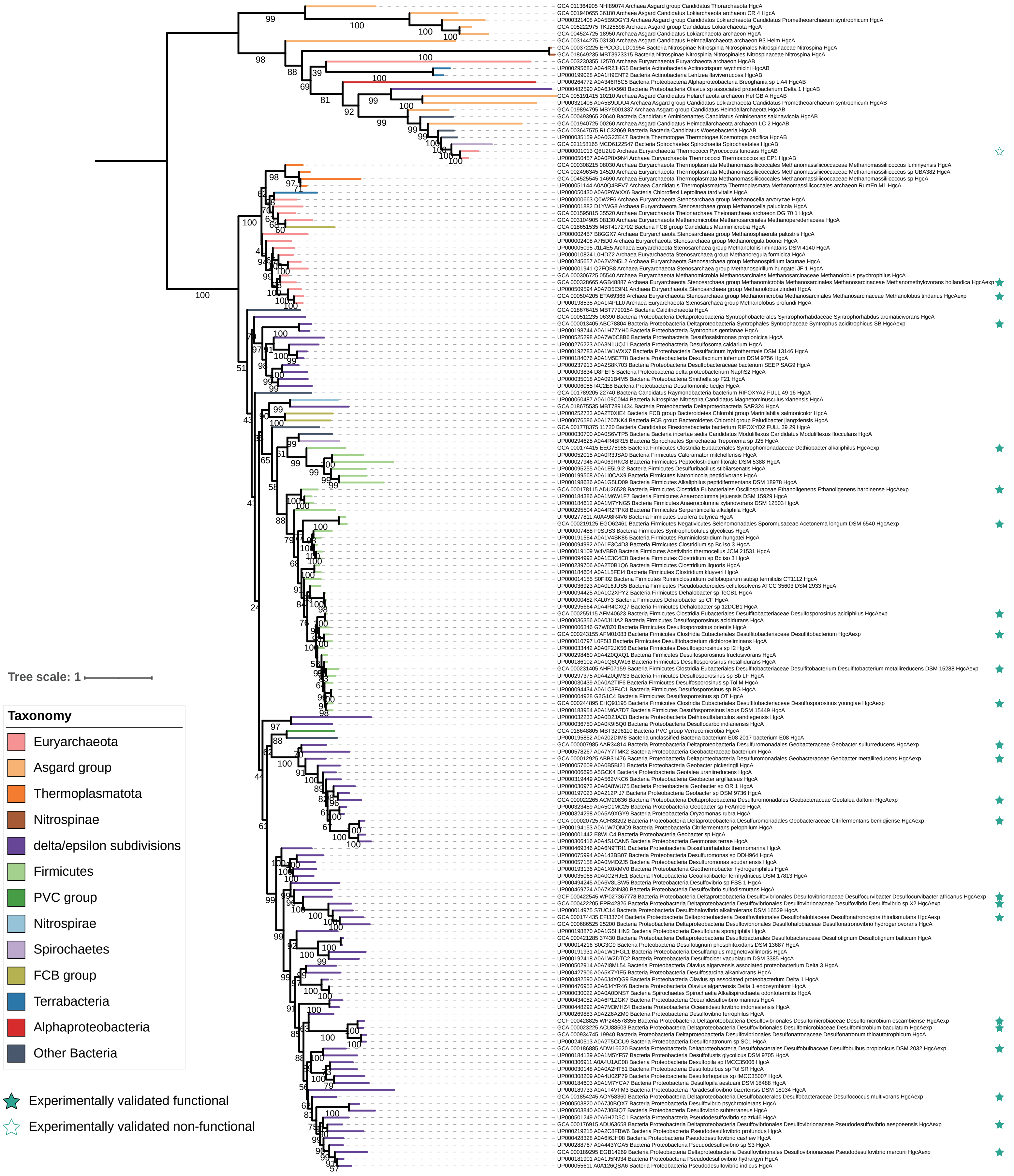


**Figure S2. Phylogenetic tree of HgcA proteins.** The tree was inferred by using the Maximum Likelihood method under LG+C60+F+G model. This analysis involved alignment of 169 amino acid sequences with a total of 493 positions. Different taxonomies are represented by different colors. Ultrabootstrap support values were calculated with 1000 replications, and ultrabootstrap values > 90% are shown by black dots at the nodes. Experimentally validated functional and non-functional HgcA sequences are labelled by solid and hollow stars. Fused-HgcAB sequences are indicated by a grey background.

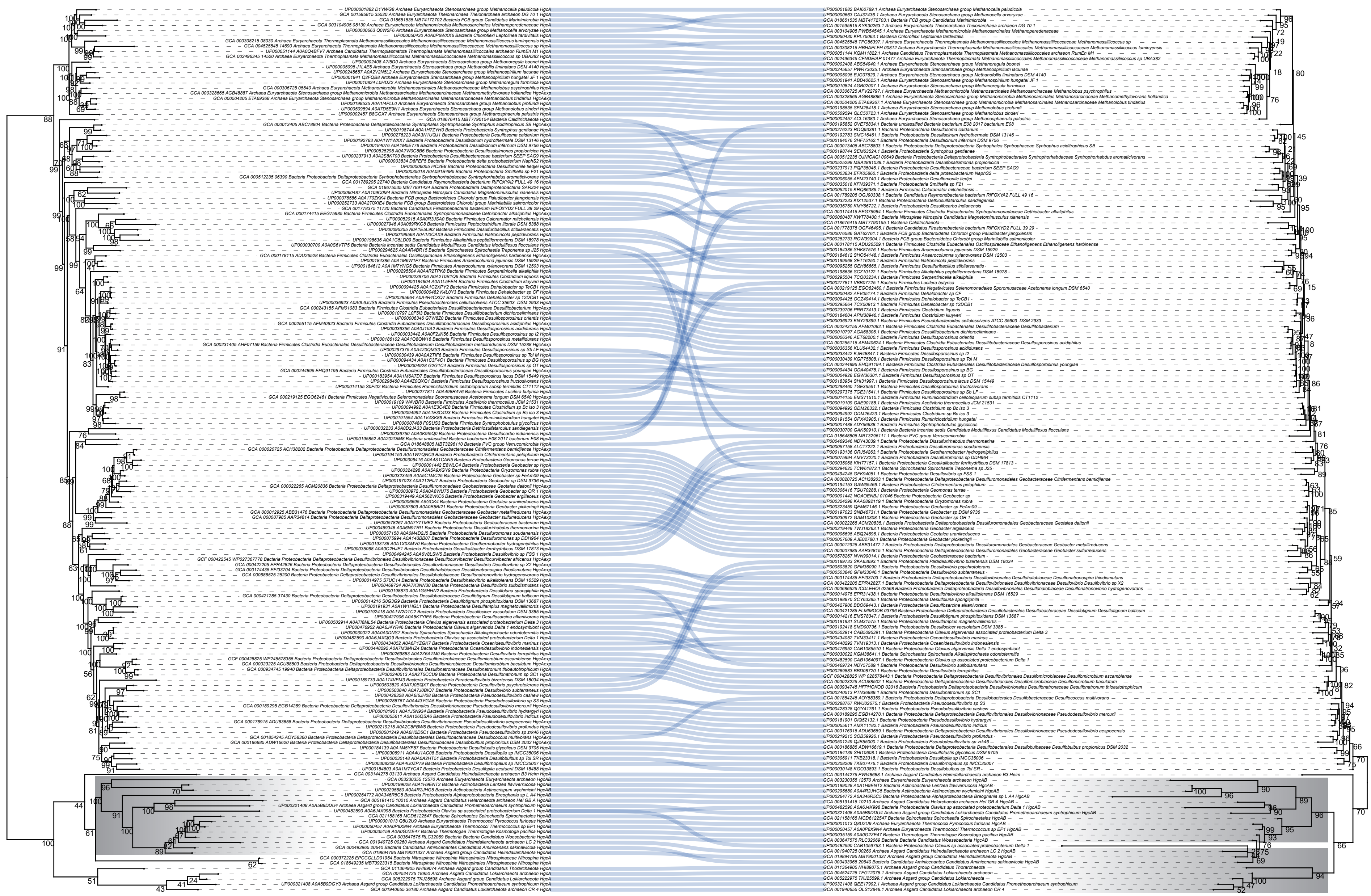

**Figure S3. Co-phylogenetic tree of HgcA and HgcB.** The phylogeny of HgcA shown on the left is the same as Figure 2. The phylogeny of HgcB shown on the right was inferred by using the Maximum Likelihood method under LG+C60+F+G model. Lines between the two trees connect HgcA (left) and HgcB (right) proteins from the same proteome, showing the phylogenetic association between the two proteins. This analysis involved alignment of 169 amino acid sequences with a total of 322 positions. Taxonomies of the HgcB sequences are represented in the same colors as shown in the HgcA phylogeny. Ultrabootstrap support values were calculated with 1000 replications, and ultrabootstrap values > 90% are shown by black dots at the nodes. Gray shaded blocks describe fused-HgcAB and 'HgcB tail' genes in the two trees, respectively. The green dots at the tip of the HgcA tree represent the corresponding *hgcA* without downstream *hgcB*, and the red triangle represents the corresponding fused-*hgcAB* with another *hgcB* downstream.

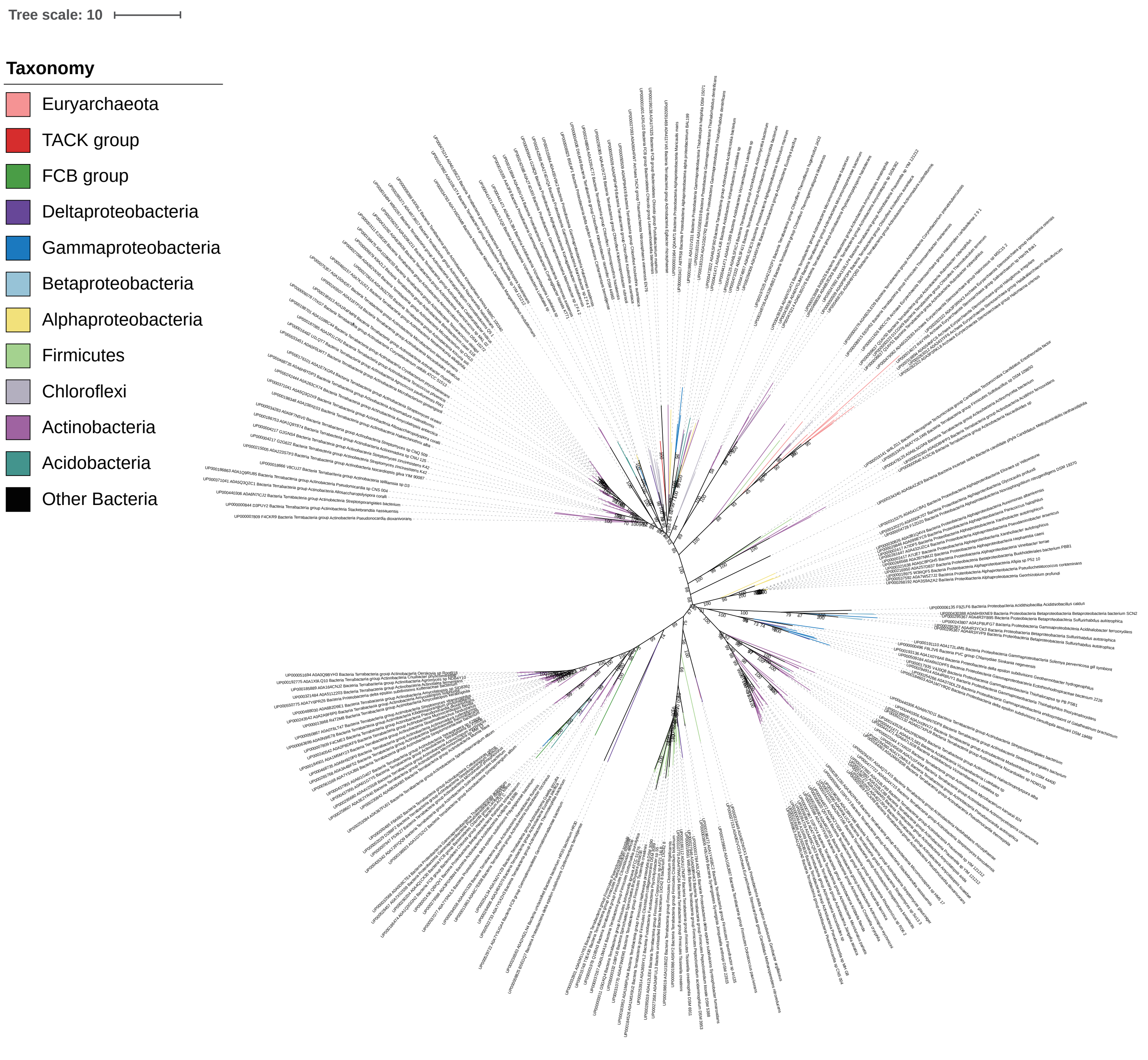


**Figure S4. Phylogenetic tree of MerB proteins.** The tree was inferred by using the Maximum Likelihood method under LG+C40+F+G model. This analysis involved alignment of 223 amino acid sequences with a total of 1448 positions. Different taxonomies are represented by different colors. Ultrabootstrap support values were calculated with 1000 replications shown at the nodes.

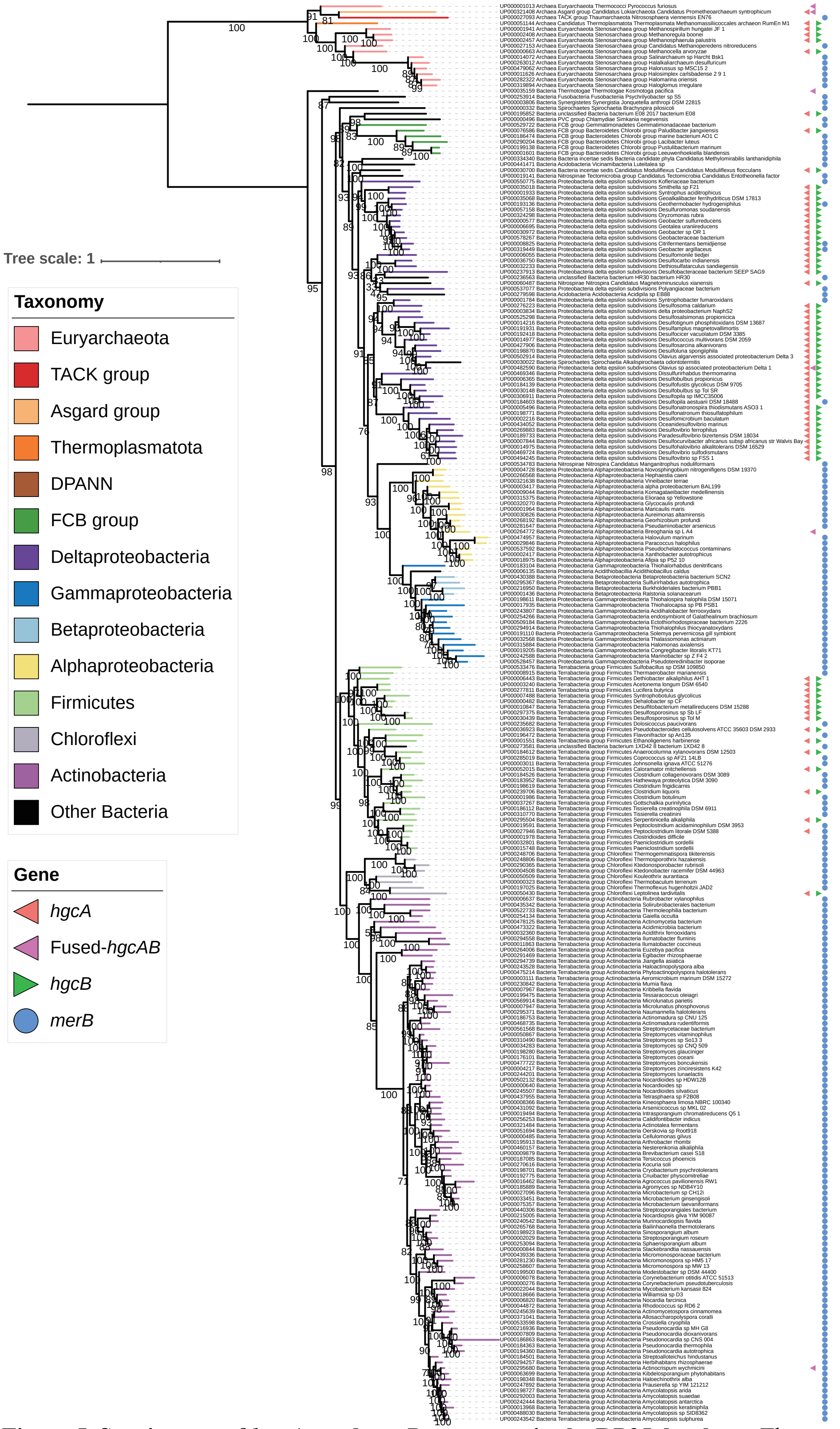


**Figure 5. Species tree of *hgcA*+ and *merB*+ genomes in the RP35 database.** The tree was inferred from a concatenate of 27 marker genes derived from Moody et al. (2022) by using the Maximum Likelihood method under LG+F+R10 model. This analysis involved alignment of 251 amino acid sequences with a total of 8024 positions. Different taxonomies are represented by different colors. The presence of genes *hgcA*/fused *hgcAB*/*hgcB*/*merB* in the genomes were represented by different symbols and colors in the outer circle. Ultrabootstrap support values were calculated with 1000 replications, and ultrabootstrap values > 90% are shown by black dots at the nodes.
